# “Modulation of apoptosis controls inhibitory interneuron number in the cortex”

**DOI:** 10.1101/134916

**Authors:** Myrto Denaxa, Guilherme Neves, Adam Rabinowitz, Sarah Kemlo, Petros Liodis, Juan Burrone, Vassilis Pachnis

## Abstract

Cortical networks are composed of excitatory projection neurons and inhibitory interneurons. Finding the right balance between the two is important for controlling overall cortical excitation and network dynamics. However, it is unclear how the correct number of cortical interneurons (CIs) is established in the mammalian forebrain. CIs are generated in excess from basal forebrain progenitors and their final numbers are adjusted via an intrinsically determined program of apoptosis that takes place during an early postnatal window. Here, we provide evidence that the extent of CI apoptosis during this critical period is plastic, cell type specific and can be reduced in a cell autonomous manner by acute increases in neuronal activity. We propose that the physiological state of the emerging neural network controls the activity levels of local CIs to modulate their numbers in a homeostatic manner.

## Introduction

The balance between excitation and inhibition (E-I balance) is essential for the generation of optimal neural circuit activity and brain function. Cortical interneurons (CIs) represent the main source of *γ*-amino butyric acid (GABA)-mediated inhibition for excitatory projection neurons (PNs) in the pallium and changes in the number or activity of CIs have been associated with neurodevelopmental and neuropsychiatric disorders, such as epilepsy, schizophrenia and autism spectrum disorders (Marin, 2012; Rubenstein and Merzenich, 2003). In contrast to cortical PNs, which are generated in the germinal zones of the dorsal telencephalon, CIs originate from progenitors in the subpallium (medial ganglionic eminence-MGE, caudal ganglionic eminence-CGE, preoptic area-POA) and following stereotypic migration routes reach the dorsal telencephalon where they integrate into the local circuits (Bartolini et al., 2013; Marin and Rubenstein, 2001; Wonders and Anderson, 2006). The disparate origin of PNs and CIs raises questions regarding the mechanisms that coordinate the size of these functionally interdependent neuronal populations of the cortex. A recent report has shown that CIs are generated in excess from basal forebrain progenitors and that BAX-dependent developmental cell death occurring over a critical postnatal period adjusts the final number of inhibitory neurons (Southwell et al., 2012). However, it is unclear whether the postnatal apoptosis of CIs is controlled by an invariable cell-intrinsic programme or can be modulated by the cellular composition and physiological state of the postnatal brain.

Systematic gene expression analysis, genetic cell-lineage tracing and phenotypic characterisation of mouse mutants have demonstrated that CI subtypes are specified by distinct region-specific transcriptional programmes operating within progenitor domains of the subpallium (Fishell and Rudy, 2011). *Lhx6* encodes a LIM homeodomain transcription factor which is specifically expressed by MGE-derived precursors and their derivative CIs expressing somatostatin (Sst) and parvalbumin (Pv). Consistent with its expression pattern, *Lhx6* mutations are characterised by a severe reduction in the number of Sst^+^ and Pv^+^ CIs, but a normal complement of GABA producing cells (Liodis et al., 2007; Zhao et al., 2008). These cellular phenotypes are associated with reduced inhibitory synaptic input on PNs, brain hyperactivity and epilepsy-like phenotypes in postnatal animals (Neves et al., 2013). Here, we have combined phenotypic analysis, genetic lineage tracing, cell transplantation and chemogenetic activation to query the specific responses of CI sub-lineages to the removal of *Lhx6* activity. We find that *Lhx6* is required to maintain the normal complement of MGE-derived CIs and that reduction of this inhibitory neuronal subpopulation in *Lhx6* mutants results in a surprising increase of Lhx6-independent CGE-derived CIs and re-balancing of the total number of CIs. The compensatory increase of CGE-derived CIs is due to a reduction in apoptosis, which can be modulated cell-autonomously by neuronal excitability during a critical postnatal period. Our results provide fundamental insight into the mechanisms that match the size of CI populations to the physiological requirements of cortical circuits and pave the way for understanding the impact of neuronal activity on cell transplantation-based therapeutic targeting.

## Results

### Loss of MGE CIs results in a compensatory increase in the number of CGE CIs

Using general (Gad1) and subtype-specific (Pv, Sst) markers for cortical inhibitory neurons, we and others have reported that mice homozygous for null mutations of *Lhx6* have a reduced number of MGE-derived Sst^+^ and Pv^+^ CI subpopulations, but the total number of GABAergic neurons in the neocortex and hippocampus remains unchanged (Liodis et al., 2007; Zhao et al., 2008). To examine the consequences of deleting *Lhx6* from specific CI lineages we used Cre-LoxP technology and targeted mutagenesis to generate a conditional allele of *Lhx6 (*Lhx6^fl^*)* in the mouse (see methods, and Fig. S1 A-B). Introduction into the *Lhx6^fl/-^* genetic background of the Cre-dependent fluorescent reporter *Rosa26-tdTomato* (tdT, Ai14, Madisen et al., 2010) allowed us to use Cre drivers for cell type-specific *Lhx6* ablation and simultaneous fate-mapping of the mutant lineages. To validate the novel *Lhx6^fl^* allele, we first used *VgatCre* (Vong et al., 2011) to delete *Lhx6* from all CI precursors. Consistent with the phenotype of *Lhx6* null mutants, the population of Pv^+^ and Sst^+^ CIs was dramatically reduced in P18 *VgatCre;Ai14;Lhx6^fl/-^* mice relative to *VgatCre;Ai14;Lhx6^fl/+^* controls (Fig. S1 C-F), while the total number of tdT^+^ cells remained unchanged (Fig. 1 A-B, G). Next, we used the *Nkx2.1Cre* transgenic driver (Kessaris et al., 2006) to track MGE-derived CIs lacking Lhx6 expression. As expected, the percentage of tdT^+^ CIs co-labelled with antibodies against Lhx6, Pv, Sst and Reelin was dramatically reduced in *Nkx2.1Cre;Ai14;Lhx6^fl/-^* mice relative to *Nkx2.1Cre;Ai14;Lhx6^fl/+^* controls (Fig. S1 G-L). Furthermore, we observed that the cortex of *Nkx2.1 Cre;Ail 4;Lhx^fl/-^* mice contained a small number of tdT^+^ cells co-expressing VIP or Sp8, markers normally associated with CGE-derived CIs (Fig. S1 M-N; see also Vogt et al., 2014). The overall number of tdT-expressing cells in the cortex of P18 *Nkx2.1Cre;Ai14;Lhx^f/-^* mice was significantly smaller relative to controls (Fig. 1 H and Fig. S2 A), suggesting that in addition to its well-established role in CI subtype specification, *Lhx6* is also required for the survival of MGE-derived CIs.

**Figure 1:**
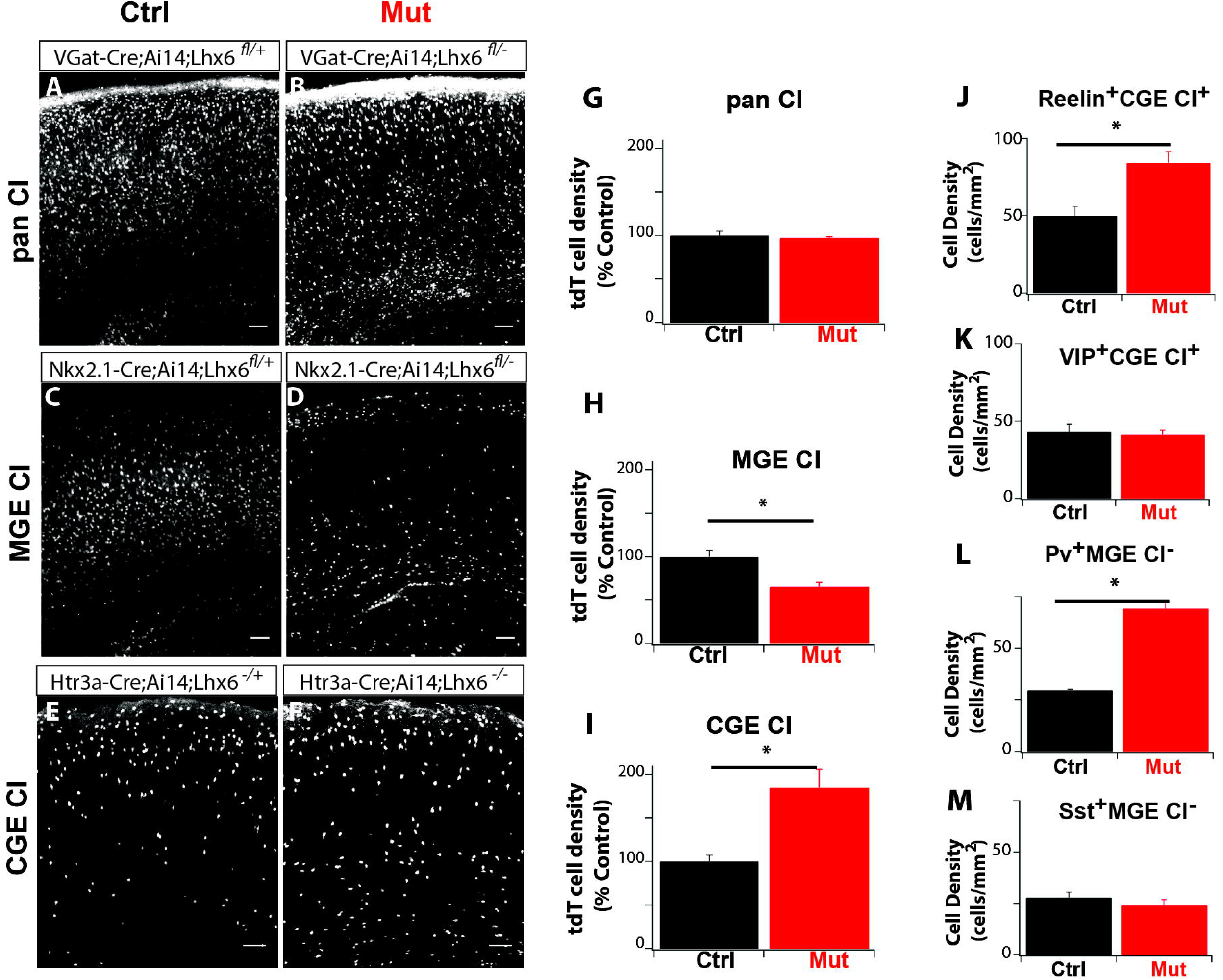
Fate mapping reveals reduced viability of *Lhx6*-deficient MGE CIs and increased survival of CGE CIs. **(A-F)** tdT^+^ (white signal) CIs in sections from the somatosensory cortex of *Lhx6* control (Ctrl, **A,C,E**) and mutant (Mut, **B,D,F**) P18 mice, fate-mapped using different Cre driver lines, as indicated in the panels. (**A,B**) The entire CI population, (**C,D**) MGE-derived, and (**E,F**) CGE-derived. (**G-I**) Quantification (normalised to the average control level) of tdT^+^ cell density in somatosensory cortices of animals represented in **A/B (G), C/D (H)** and **E/F (I)**. **(J-M)** Quantification of cell densities of CGE-derived Reelin^+^ **(J)**, CGE-derived VIP^+^ **(K)**, non-MGE-derived Pv^+^ **(L)** and non-MGE-derived Sst^+^ **(M)**. Similar results were observed in the Motor Cortex (Fig. S2). Data expressed as mean ± SEM. Statistical significance evaluated using Student’s t-test, * denotes p < 0.05, ** p<0.005.

The reduced number of tdT^+^ MGE CIs observed in the cortex after deleting *Lhx6 (Nkx2.1Cre;Ai14;Lhx6^fl/-^* mice, Figs. 1 C-D, H, S2A), compared to the near normal number of GABAergic interneurons in the cortex of either *Lhx6* null mice (Liodis et al., 2007; Zhao et al., 2008) or mice where *Lhx6* is deleted from all CIs (*VgatCre;Ai14;Lhx6^fl/-^*; Fig. 1 G), suggests that non-MGE-derived CI lineages compensate for the specific loss of MGE-derived CIs. To examine this possibility we fate-mapped CGE-derived CIs in *Lhx6^-/-^* mice using the CGE-specific Cre driver *Htr3αCre* (www.gensat.org; see methods) and the Ai14 (tdT) reporter. We observed an increased representation of CGE-derived tdT^+^ cells in the cortex of *Lhx6* mutant mice *(Htr3aCre;Ai14;Lhx6^-/-^)* relative to controls *(Htr3aCre;Ai14;Lhx6^+/-^)* (Figs. 1 E-F, I, S2 D). Therefore, the size of the CGE-derived CI population in the mammalian cortex is not pre-determined but can be modulated to compensate for the loss of MGE-derived interneurons in the cortex.

The majority of CGE-derived CIs can be accounted for by two functionally and molecularly distinct sub-populations marked by VIP or Reelin (Fishell and Rudy, 2011; Kepecs and Fishell, 2014; Lee et al., 2010). Immunostaining of *Lhx6* mutant brain sections in CGE labelled CI mice *(Htr3aCre;Ai14;Lhx6^-/-^)* using these subtype markers showed that only the Reelin^+^ subset increased in number while the VIP^+^ subpopulation remained unchanged (Figs. 1 J-K, S2 E-F). Interestingly, ablation of *Lhx6* from MGE lineages also resulted in altered representation of CI subtypes originating outside the ganglionic eminences (Gelman et al., 2011). Thus, in *Nkx2.1Cre;Ai14;Lhx6^fl/-^* mice the number of tdT^-^Pv^+^ interneurons (which are likely to originate from the preoptic area-POA, see Fig. S3) increased relative to controls (*Nkx2.1Cre;Ai14;Lhx6^fl/+^)* while the number of tdT^-^Sst^+^ CIs remained unchanged (Figs. 1 L-M and S2 B-C). These findings suggest that the compensatory responses of CIs to *Lhx6* ablation are subtype-specific and occur across different lineages.

### Reciprocal changes in apoptosis of CGE and MGE CIs in Lhx6 mutants

The increased representation of CIs originating outside the MGE in the cortex of *Lhx6*-deficient mice could result from enhanced proliferation of their progenitors or reduced neuronal cell death during the critical postnatal window of apoptosis (Southwell et al., 2012; Yamaguchi and Miura, 2015). To distinguish between these possibilities we first compared the number of proliferating progenitors (identified by pH3 immunostaining and EdU uptake) within the ganglionic eminences of *Lhx6* mutant and control E14.5-16.5 embryos. The estimated number of pH3^+^ and EdU^+^ progenitors was similar between the two genotypes, suggesting that the mechanism(s) responsible for the increased representation of CGE-derived CIs in *Lhx6* mutants operates on post-mitotic interneuron precursors at later developmental stages (Fig. S4 A-H). In agreement with this hypothesis, there was no difference in the number of cells expressing Sp8 (a transcription factor expressed by non-MGE derived CIs, Ma et al., 2012) in E16 control and mutant cortices (Fig. S4 I-L). In further support of this idea, the number of fate-mapped MGE-and CGE-derived CIs at P2 (a developmental stage that follows the completion of CI tangential migration but precedes the onset of apoptosis, Miyoshi and Fishell, 2011; Southwell et al., 2012) was similar between control and *Lhx6* mutant mice (Fig. 2 A-D and I). In contrast, 5 days later (at P7) we observed a decreased representation of MGE-derived CIs and a reciprocal increase of their CGE-derived counterparts (Fig. 2 E-H and I). The changes in the size of MGE-and CGE-derived CIs populations observed at P7 foreshadow those observed at P18 (Figs. 1 H, I and 2 I) and together suggest lineage-specific modulation of the apoptotic programmes of CIs by the *Lhx6* mutation.

**Figure 2:**
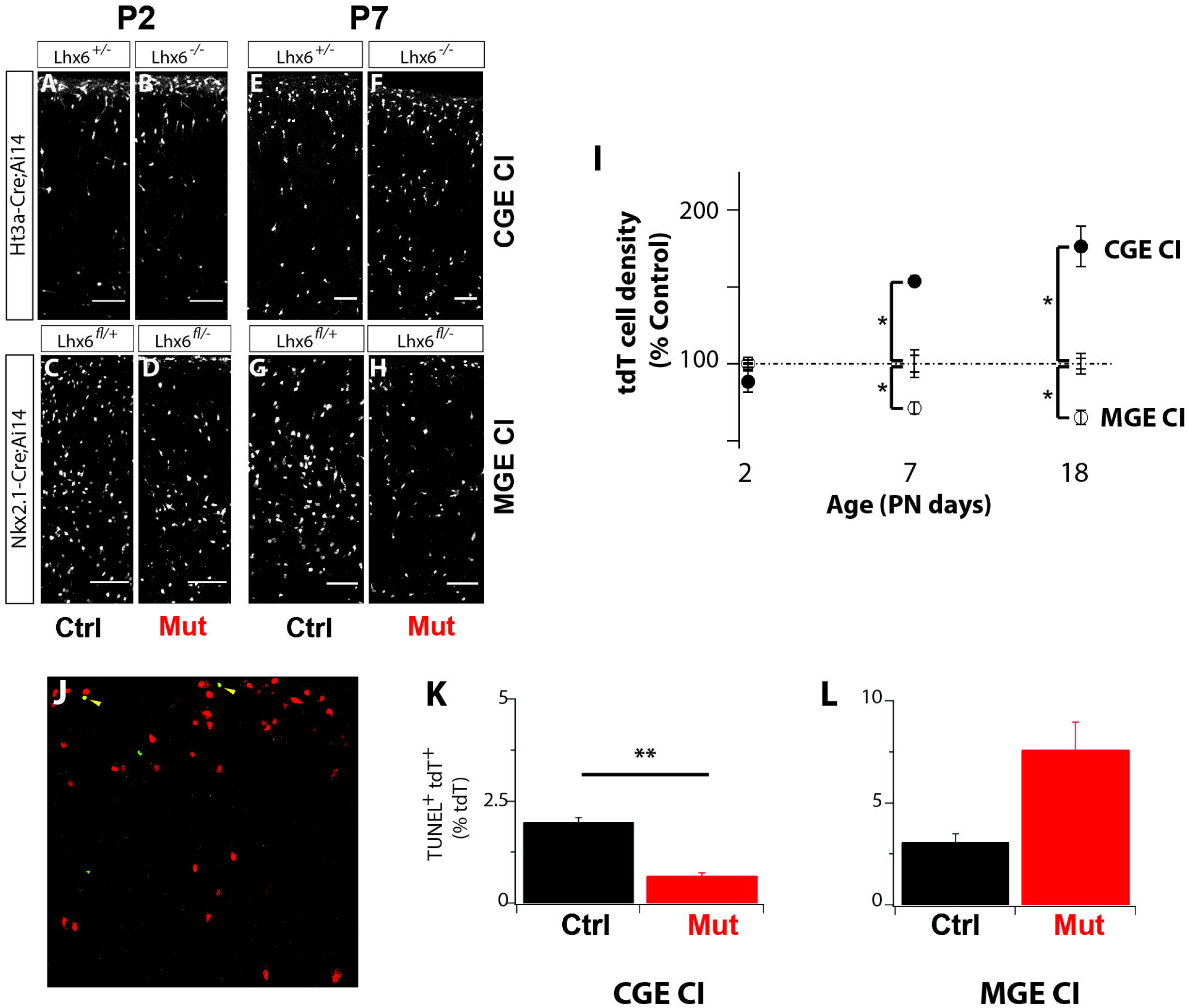
Reduced apoptosis of CGE CIs compensates for the loss of *Lhx6*-deficient MGE counterparts. Changes in the number of CI subtypes in *Lhx6* control and mutant mice during early post-natal development. (**A-H**) tdT-expressing CIs in cortical sections from P2 (**A-D**) and P7 (**E-H**) *Lhx6* control (**A,C,E,G**) and mutant (**B,D,F,H**) mice. tdT expression identifies CGE-derived (**A,B,E,F**) and MGE-derived CIs (**C,D,G,H**) respectively. (**I**) Quantification of CGE (close circles) and MGE (open circles) CI density in the whole cortex of *Lhx6* mutant mice relative to *Lhx6* controls (dotted line) at P2, P7 and P18. (**J**) Representative cortical section from P7 control mice showing MGE derived CIs (tdT^+^ in red) and TUNEL+ (green) cells. Yellow arrowheads indicate TUNEL^+^ MGE derived CIs. (**K,L**) Quantification of the fraction of CGE (**K**) and MGE (**L**) CIs undergoing apoptosis (tdT^+^TUNEL^+^) in the cortex of *Lhx6* control and mutant P7 mice.

To confirm this, we used TUNEL staining to quantify the extent of developmental cell death of MGE-and CGE-derived CIs at P7 *Lhx6* mutants, when expression of apoptotic markers in these cells is at its highest (Fig. 2 J; Southwell et al., 2012). Although CGE-derived CIs do not express *Lhx6*, the number of tdT^+^TUNEL^+^ double positive CIs in the cortex of P7 *Htr3aCre;Ai14;Lhx6^-/-^* mice was significantly reduced relative to control littermates *(Htr3aCre;Ai14;Lhx6^+/-^;* Fig. 2 K). Similar analysis in the cortex of *Nkx2.1Cre;Ai14;Lhx6^fl/-^* P7 mice revealed a clear trend for an increase in the number of tdT^+^TUNEL^+^ cells relative to control animals *(Nkx2.1Cre;Ai14;Lhx6^fl/-^;* Fig. 2 L). We suggest that ablation of *Lhx6* during embryogenesis results in increased representation of non-MGE-derived CIs via non-cell autonomous effects on their developmental cell death programmes.

### Enhanced survival of wild-type CIs grafted into the Lhx6 mutant cortex

To explore the possibility that the cortical microenvironment of *Lhx6* mutants can rescue CIs from apoptosis, we transplanted GFP^+^ CI precursors isolated from the basal forebrain of GAD-GFP E14.5 embryos (Tamamaki et al., 2003) into the pallium of neonatal (P0-P1) *Lhx6^-/-^* pups and their control *(Lhx6^+/-^)* littermates (Fig. 3 A). Multiple morphologically mature GFP^+^ CIs were observed throughout the cortex at P16, in both control and mutant animals (Fig. 3 B-F). However, a subset of grafted CIs in mutant cortices showed striking morphologies, with consistently larger somas and dendritic arbours, a finding that mirrored the unusually large size of endogenous POA-derived Pv^+^ CIs observed in *Lhx6* mutant mice (Fig. S5).

**Figure 3:**
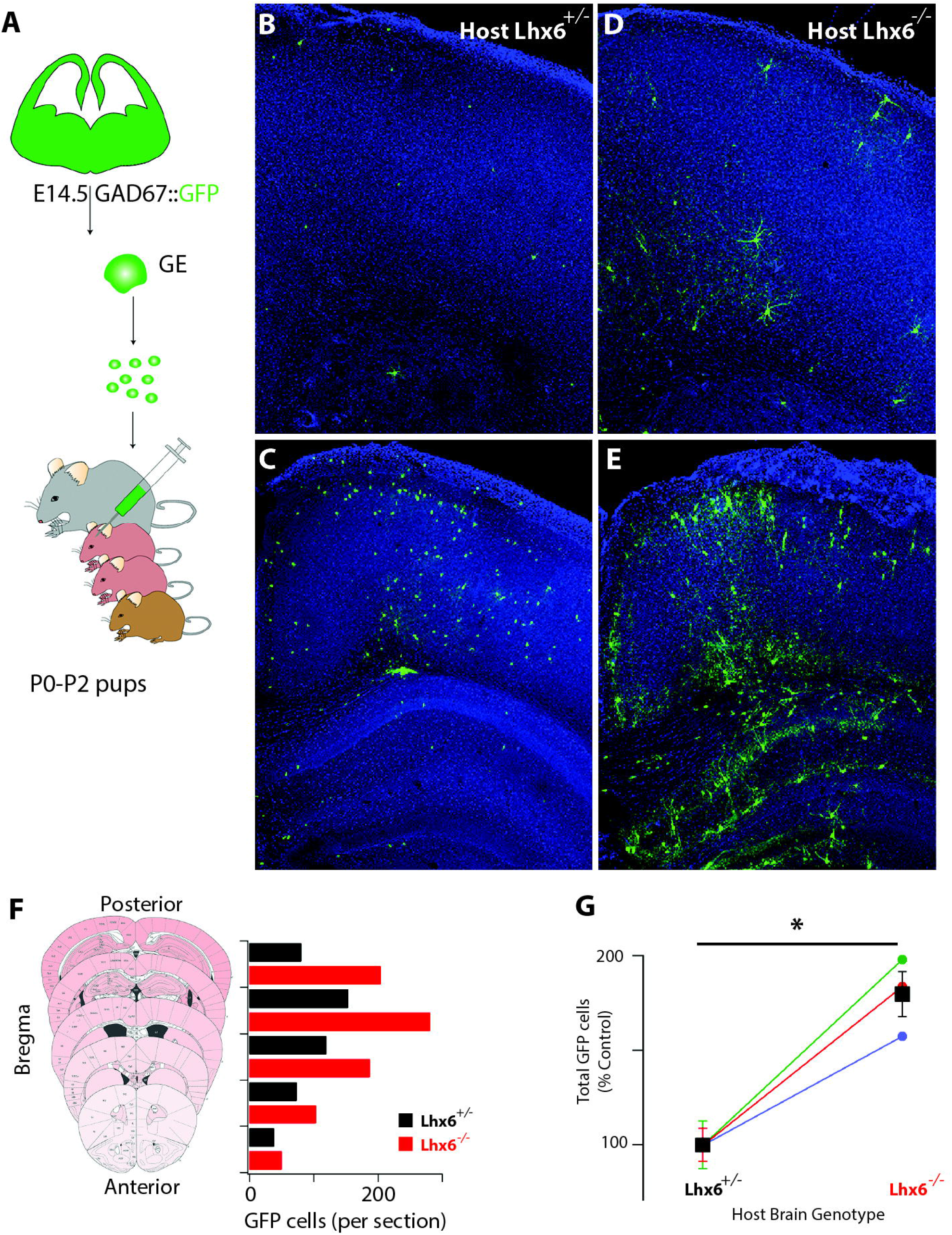
Cellular microenvironment of **Lhx6** mutant forebrain promotes grafted CI progenitor survival. **(A)** Schematic representation of the CI progenitor transplantation into the cortex of P0-P2 pups. **(B-E)** Coronal brain sections of Lhx6^+/-^ **(B,C)** and Lhx6^-/-^ **(D,E)** P16 mice transplanted at P0-P2 with wild-type GFP^+^ CI progenitors. Sections shown in **B,D** are more anterior to those shown in **C,E**. **(F)** Quantification of surviving GFP^+^ CIs in the cortex of one *Lhx6^+/-^* (black bars) and *Lhx6^-/-^* (red bars) littermate pair of P16 mice grafted with CI precursors at P0-P2, at different rostro-caudal levels. **(G)** Quantification of surviving GFP^+^ CIs in the cortex of *Lhx6^+/-^* and *Lhx6^-/-^* P16 mice. Values for the mutants (*Lhx6*^-/-^) (circles) are normalized to the average value in control littermates *(Lhx6^+/-^;* coloured bars represent SEM for individual grafting experiments). The number of GFP^+^ cells found in *Lhx6^-/-^* was 181 ± 13% higher in comparison to control littermates (p=0.02 one sample t-test - mean 100 - n= 3 *Lhx6^-/-^*, 10 *Lhx6^+/-^*; minimum of 500 cells counted per brain).

The majority of grafted CI precursors are eliminated by BAX-dependent apoptosis within 2 weeks of transplantation (Southwell et al., 2012), a feature that recapitulates the time-line of programmed cell death of endogenous CIs. Consistent with this idea, we found that the number of transplanted GFP^+^ CIs in the cortex of P16 *Lhx6^-/-^* mice was higher relative to that in *Lhx6^+/-^* littermate controls (181 ± 13% of control; p=0.02, n=3 litters; Fig. 3 G), confirming the notion that the microenvironment of the host brain can modulate the survival of interneurons in the cortex.

### Transcriptomic analysis of Lhx6-deficient brains/lineaşes

To provide insight into the mechanisms by which the cortical microenvironment of *Lhx6* mutants controls the survival of grafted CI progenitors, we used RNA sequencing (RNAseq) to compare the global transcriptome of the forebrain dissected from control *(Lhx6^+/-^)* and mutant *(Lhx6^-/-^)* mice at P15, a stage at which apoptosis of grafted CIs is at its highest. Differential gene expression analysis identified 1707 genes that were significantly up-regulated (977) or down-regulated (730) (Fig. 4 A) in mutant relative to control littermates. In addition to *Lhx6* (which was absent from mutant samples), *Sst* and *Pv* were among the top down-regulated genes in *Lhx6* mutants, in agreement with our immunocytochemistry data (Fig. S1). Interestingly, inspection of the list of up-regulated genes identified several genes - including *Bdnf* (Hartmann et al., 2001), *Npas4* (Bloodgood et al., 2013), *Fosb* (Eagle et al., 2015) and *Npy* (Gall et al., 1990) - whose expression is induced by neuronal activity. Gene set enrichment analysis demonstrated that genes previously shown to be up-regulated in pyramidal neurons following chronic increases in activity (through inhibition of ionotropic GABA receptors, Yu et al., 2015), were similarly enriched in *Lhx6*-deficient samples (false discovery rate<10^-59^). In fact, hierarchical clustering based on the expression of the top 25 genes up-regulated by chronic activity clearly distinguished between control and mutant samples (Fig. 4 B). Finally, the expression of the activity-dependent gene *cfos* (Cohen and Greenberg, 2008) was highly up-regulated in the cortex of *Lhx6* mutants (Fig. 4 C-D). These transcriptomic results provide a molecular confirmation of increased network activity in the cortex, as would be expected for brains where the development of MGE-derived CIs is compromised (Batista-Brito et al., 2009; Neves et al., 2013).

**Figure 4:**
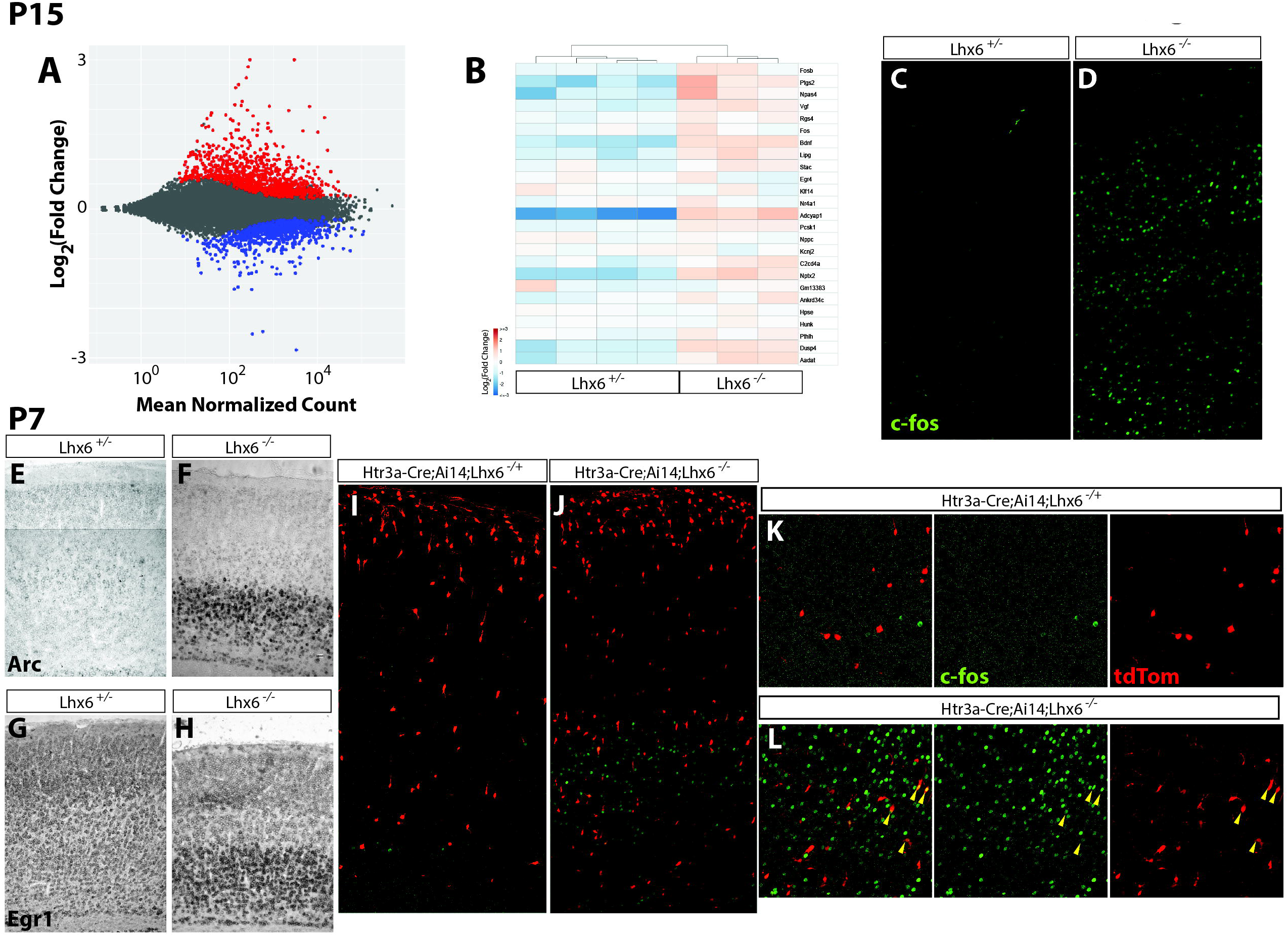
Molecular analysis of *Lhx6* mutant forebrains reveal widespread up-regulation of activity-dependent genes. **(A)** MA plot summarizing the results of the differential expression analysis between *Lhx6^+/-^* and *Lhx6^-/-^* P15 forebrains. Significantly upregulated genes are shown in red (977 genes) while significantly down-regulated genes are shown in blue (730 genes). Significance was set as a false discovery rate of <= 0.05. **(B)** Hierarchical clustering of *Lhx6^+/-^* and *Lhx6^-/-^* forebrain samples. Clusters were generated using the expression levels of the 25 most significantly upregulated genes in dissociated hippocampal cultures upon chronic treatment with bicucullin (Yu et al., 2015). Genes upregulated upon bicucullin treatment are similarly upregulated upon deletion of *Lhx6*. **(C,D)** Coronal sections from the somatosensory cortex of *Lhx6^+/-^* **(C)** and *Lhx6^-/-^* **(D)** P15 mice immunostained for cfos (green). **(E-H)** *In situ* hybridization of somatosensory cortex sections from *Lhx6^+/-^* **(E,G)** and *Lhx6^-/-^* **(F,H)** P7 mice with either *Arc* **(E,F)** or *Egr1* **(G,H)** riboprobes shows immediately-early gene upregulation in the bottom layers of mutant brains. **(I-J)** Coronal sections of the somatosensory cortex of *Htr3a-Cre;Ai14;Lhx6^+/-^* **(I)** and *Htr3a-Cre;Ai14;Lhx6^-/-^* **(J)** P7 mice immunostained for cfos (green) reveal CGE CIs present in mutant cortices up-regulate cfos expression. **(K-L)** show higher magnification images of **(I-J)** respectively, arrowheads show examples of cfos expressing CGE derived tdT^+^ cells.

Next, we compared the expression of neuronal activity markers between control and *Lhx6*-deficient cortices at P7, a postnatal stage characterised by the highest rate of interneuron cell death (Southwell et al., 2012). This analysis showed a dramatic upregulation of a number of immediate early genes including the activity regulated cytoskeleton-associated protein *Arc* (Tzingounis and Nicoll, 2006), the early growth response protein *Egr1* (French et al., 2001) and *cfos* in the cortex of *Lhx6*-deficient mice relative to controls (Fig. 4 E-H). Interestingly, overexpression of the activity-dependent markers was observed mostly in the bottom layers of the cortex, which also showed the highest increase in the number of CGE-derived CIs (Figs. 1 E-F, 2 E-F; the fraction of CGE derived tdT^+^ cells present in the bottom cortical layers increases from 15 ± 2 % in controls to 47 ± 2 % in mutants, P18 cortices, p=0.02). Together, our gene expression analysis demonstrates a correlation between increased immediate-early gene expression, which is reflective of enhanced network activity, and improved survival of CIs in the cortex of *Lhx6* mutant mice.

Next, we compared the expression of immediate-early gene markers specifically in CGE-derived CIs labelled with tdT in *Htr3aCre;Ai14;Lhx6^-/-^* versus *Htr3aCre;Ai14;Lhx6^+/-^* mice at P7. First, *cfos* immunostaining showed that the number of cFos^+^tdT^+^ neurons (yellow arrowheads in Fig. 4 L) was increased in *Lhx6*-deficient cortex relative to controls, suggesting increases in the activity of CGE CIs (Fig. 4 I-L). To further characterise such changes, we employed RT^2^ profiler PCR array technology (see methods) to compare the expression of a panel of known activity-associated genes in CGE-derived CIs isolated by flow cytometry from the brain of P7 *Htr3aCre;Ai14;Lhx6^-/-^* and *Htr3aCre;Ai14;Lhx6^+/-^* mice (Table S2). Among the genes upregulated (>1.5 fold change) in CGE CIs from *Lhx6*-deficient brains, were a number of genes associated with increased activity levels, including two members of the EGR family *(Egr2* and *Egr3;* DeSteno and Schmauss, 2008; Li et al., 2007), the neurotrophic factor *bdnf* (Hartmann et al., 2001), as well as genes implicated in growth factor signalling, such as the insulin-like growth factor 1 (*Igf1*; Mardinly et al., 2016) and the nerve growth factor receptor *(Ngfr,* Meeker and Williams, 2015). These factors and their receptors play a crucial role in the control of neuronal numbers and dendritic growth. Together, these findings identify increased activity of CIs as a potential mechanism that drives their enhanced survival in hyperactive cortical networks.

### Cell autonomous increase in the activity of CIs enhances survival

To directly test whether CI survival is regulated by neuronal activity in a cell autonomous manner, we transplanted CI precursors expressing Designer Receptors Exclusively Activated by Designer Drugs (DREADDs) and modulated their activity by administering appropriate ligands (Urban and Roth, 2015). Specifically, the MGE of E14.5 embryos was co-electroporated with a bi-cistronic expression vector encoding the hM3D(Gq) DREADD and Red Fluorescent Protein (RFP) and a control plasmid encoding GFP. Transfected CIs were mechanically dissociated and the resulting cell suspension grafted in the cortex of P0-P1 wild type mice. Since only a fraction of electroporated (GFP^+^) neurons co-expressed hM3D(Gq) (RFP^+^) (Fig. 5 A-E), the GFP^+^RFP^-^ population served as an internal control for the effect of DREADD ligands. Indeed, administration of the DREADD ligand clozapine-N-oxide (CNO) selectively increased the activity of transfected GFP^+^RFP^+^ cells (Fig. S6). Importantly, CNO treatment (administered twice daily from P14-P17) resulted in an increase in the fraction of GFP^+^RFP^+^ (yellow arrowheads) relative to GFP^+^RFP^-^ (white arrows) cells when compared to vehicle administered littermates (Fig. 5 F-J), suggesting that enhanced activity is sufficient to protect CIs from programmed cell death in an otherwise normal brain. Our data provide evidence that neuronal activity modulates the number of CIs in the cortex in a cell autonomous manner.

**Figure 5:**
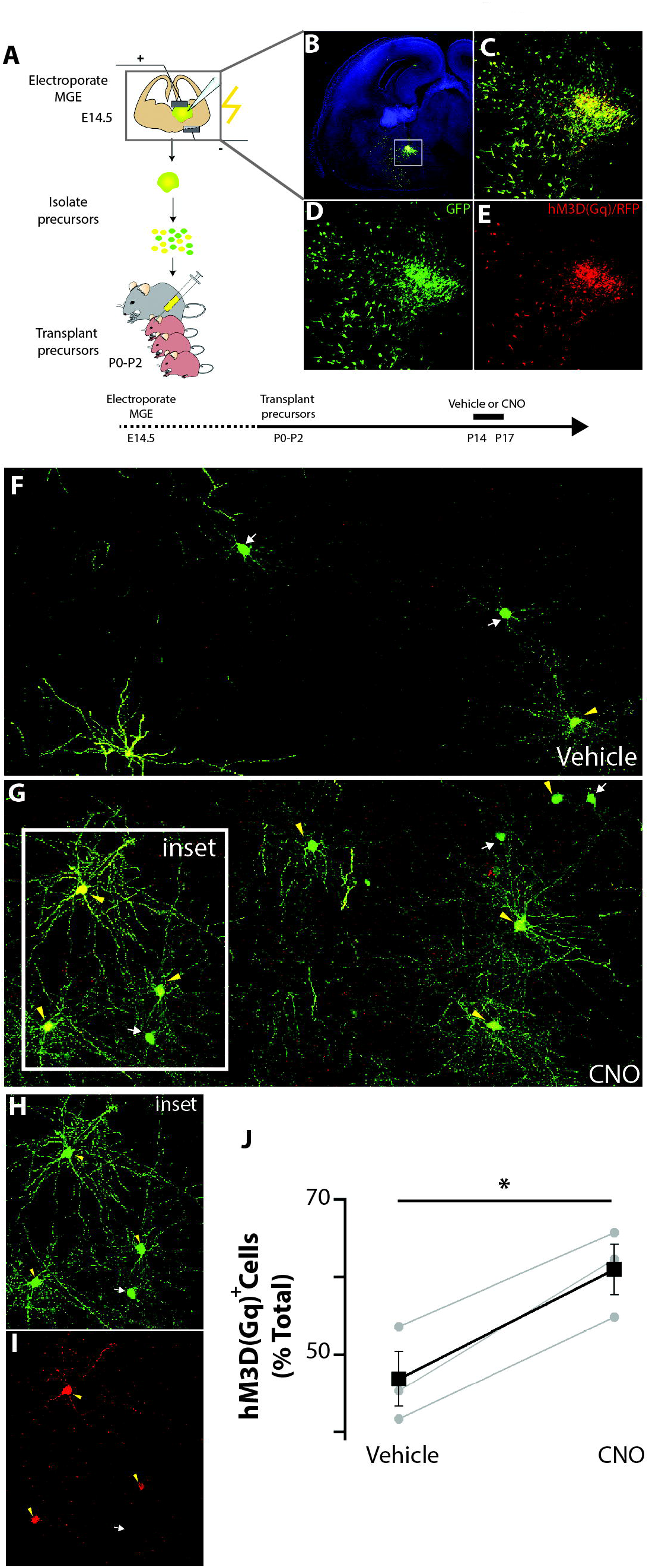
Cell autonomous depolarization of CIs enhances their survival. **(A)** Schematic representation of brain acute slice electroporation, grafting and vehicle/CNO administration protocol. Drug administration was targeted to coincide with the peak of apoptosis of transplanted CI progenitors. **(B)** Coronal section from an E14.5 embryo brain transfected with CAG:IRES:GFP (pGFP) and CAG:hM3D(Gq):IRES:RFP (pDREADRFP) plasmids and cultured for 12 h. Boxed area is magnified to show expression of both fluorescent reporters **(C)**, GFP only **(D)** and RFP only **(E)**. **(F-I)** Representative sections from the somatosensory cortex of P17 mice grafted at P0-P2 with CI precursors transfected with pGFP and pDREADDRFP plasmids and injected with either vehicle **(F)** or CNO **(G)**. Yellow arrowheads identify cells expressing both plasmids, while green arrows show cells expressing GFP only **(H,I)** Boxed region in G magnified to reveal expression of GFP **(H)** and RFP **(I)**. **(J)** Quantification of RFP^+^ cells found in the forebrain of P17 mice transplanted at P0-P2 (normalized to the total GFP^+^ population). (RFP^+^ - vehicle = 47 ± 3 %; CNO = 61 ± 3 %, p=0.01 Student’s paired samples t-test, n= 3 vehicle, 3 CNO, minimum of 150 cells counted per brain).

## Discussion

Distinct physiological mechanisms, collectively referred to as homeostatic plasticity, operate in the nervous system to maintain or restore the balance between excitation and inhibition, even after considerable disruption of network dynamics (Turrigiano, 2012). For such “acute” mechanisms to be effective, it is essential that all physiologically relevant cellular compartments achieve a critical size and composition during development. How the output of developmental programmes that specify the number and subtypes of neurons matches the functional requirements of mature neuronal circuits remains unclear. Here, we provide evidence that modulation of programmed cell death during a critical early postnatal period is a regulatory mechanism that controls in a homeostatic manner the number of GABAergic interneurons in the mammalian cortex. Our experiments highlight a critical interplay between the physiological state of the network and its cellular units and suggest a feedback mechanism that fine-tunes the size of the CI population to stabilise brain activity.

Early stages of neural development are often characterised by large-scale proliferative expansion of progenitors and the generation of surplus number of neurons, which are eliminated at later stages by apoptosis in order to meet the physiological requirements of the system. For example, the size of motor neuron and sympathetic neuron pools is largely determined during development by the availability of limiting amounts of retrograde prosurvival signals supplied by appropriate peripheral targets (Davies, 2003; Oppenheim, 1991). However, the neurotrophic factor paradigm cannot explain adequately the regulation of apoptosis in most regions of the CNS where alternative pathways have been implicated (Dekkers et al., 2013). Earlier *in vivo* and *in vitro* studies have demonstrated that survival of cortical PNs is enhanced by network activity and that NMDA receptor-mediated changes in synaptic activity modulate rates of apoptosis (Blanquie et al., 2017; Ikonomidou et al., 1999; reviewed by Bell and Hardingham, 2011) (Bell and Hardingham, 2011) (Bell and Hardingham, 2011) (Bell and Hardingham, 2011) (Bell and Hardingham, 2011). In addition, apoptosis of adult generated neurons, such as olfactory bulb interneurons and dentate gyrus granule cells, can be dramatically influenced by the activity of the mature networks they integrate into (Bovetti et al., 2009; Corotto et al., 1994; Mu et al., 2015; Petreanu and Alvarez-Buylla, 2002; Rochefort et al., 2002; Tashiro et al., 2006), probably through a cell autonomous mechanism (Lin et al., 2010). Contrasting these studies, Southwell and colleagues demonstrated recently that in rodents a large fraction of CIs (~40%) generated in the basal forebrain during development are eliminated during a critical early postnatal period by a program of apoptosis that is intrinsic to this cell lineage (Southwell et al., 2012). Our genetic lineage tracing experiments in *Lhx6* mutants argue against a rigid and intrinsically determined programme of apoptosis of CI progenitors but rather suggest a considerable degree of developmental plasticity driven ultimately by the physiological state of the network. Although the upstream physiological parameters that characterise an “anti-apoptotic” network are currently unclear, we provide evidence that CIs monitor the activity of their local microenvironment and adjust the level of apoptosis in a cell-autonomous manner. Although we do not provide the network mechanism that controls CI apoptosis, the overall activity of local circuits provides a likely candidate. This view is supported by transplantation studies demonstrating the preferential survival of either wild-type CI progenitors grafted into the hyperactive cortex of *Lhx6*-deficient animals (Fig. 3) or by chemogenetically-activated CIs grafted into the cortex of wild type animals (Fig. 5). Together with the modulation of endogenous CI numbers in the *Lhx6* mutant cortex, our findings suggest an overarching mechanism for the control CI number in the pallium during development, whereby inhibitory interneurons monitor the activity of their local environment and adjust the level of apoptosis in a cell-autonomous manner. We hypothesise that these mechanisms act only in interneurons that have the same cellular age (Southwell et al., 2012), and therefore the same maturation state, and thus compete among themselves for their integration into cortical networks. However, the precise intracellular cascades that link the properties of the local network with the activation of CIs and their increased survival remain unclear. Interestingly, the enhanced apoptosis of immature CIs observed in response to pharmacological inhibition of NMDA receptors (Roux et al., 2015) suggests that excitatory glutamatergic neurotransmission may play a role in this regulatory pro-survival response. In addition, our own transcriptomic analysis has identified a set of candidate genes that may help uncover the mechanisms behind activity regulated apoptosis and should be investigated further.

Other studies have also shown apparent compensatory forms of plasticity in response to the loss of CI subtypes. Deletion of the *Lhx6*-dependent effector gene *Sox6* in postmitotic immature interneurons was associated with a dramatic decrease in the number of Pv^+^ and Sst^+^ interneurons, without an obvious change in the total number of CIs (Azim et al., 2009; Batista-Brito et al., 2009). Although an increase of NPY^+^ interneurons was reported in these studies, the mechanisms that maintain the total number of CIs in *Sox6* mutants remain unclear. Also, conditional inactivation of the orphan nuclear receptor *Nrf1 (COUP-TFI)* in interneuron progenitors resulted in decreased number of CR^+^ and VIP^+^ CIs and a concomitant increase of PV-and NPY-expressing subtypes, without affecting the total number of GABAergic interneurons in the cortex (Lodato et al., 2011). Although the authors proposed that this compensation is due to enhanced CI progenitor proliferation, it is possible that changes in apoptosis may also have contributed to this phenotype. Our data shows that only certain subtypes of CIs regulate their rate of apoptosis. For example, among the *Lhx6*-independent CGE-generated CIs only Reelin^+^ neurons increase in number following *Lhx6* deletion (Figs. 1 and S2). The subtype-specific response of CIs to activity is not solely restricted to apoptosis, as recent studies have shown that silencing CGE-derived interneurons results in defects in radial migration, cell morphology and synaptic development of Reelin^+^ but not VIP^+^ CIs (De Marco Garcia et al., 2011; De Marco Garcia et al., 2015). The heterogeneous response of CI subtypes to ongoing network activity during development could have important implication for the establishment of the repertoire and connectivity pattern of local circuits in the cortex. Specifically, regional differences in activity patterns in the developing brain (Allene et al., 2008; Allene et al., 2012; Blanquie et al., 2017; Khazipov et al., 2004) may be responsible for the variability in CI number or subtype found in the mature cortex. Exploring the relationship between brain activity and CI repertoire will provide important insight into how local circuits are assembled.

Our results provide evidence for a simple mechanism that controls the number of inhibitory interneurons in the cortex. We propose that the temporal overlap between developmental programmes that dictate the size (and thus the functional output) of the CI complement and the emerging activity of cortical networks allows for the engagement and cross-regulation of the two processes until an optimal activity set point is attained. Several preclinical models of CI-based cell therapies have been established for the treatment of epileptic seizures (Alvarez Dolado and Broccoli, 2011; Southwell et al., 2014; Tyson and Anderson, 2014). Our present data argue that increased activity levels in the host brain, typically observed in epileptic encephalopathy mouse models (Batista-Brito et al., 2009; Hedrich et al., 2014), or increased activity in transplanted CIs, will provide favourable conditions for the survival of grafted CIs progenitors. Characterizing the pro-survival patterns of neuronal activity and identifying the CI subtypes best suited for transplantation may improve the effectiveness of these nascent therapies.

## Material and Methods

### Lhx6 conditional mice and genotyping

A conditional *Lhx6* allele was generated via homologous recombination using a targeting construct in which loxP sites were placed in non-coding regions just 5’ to coding exon 1b, and 3’ to coding exon 3 (Fig. S1 A). The 5’ homology is made up from a 3.1 kb *XbaI-Sac*I fragment containing the 5’ upstream region and the first exon (1a) of *Lhx6,* whereas the 3’ homology is made up from a 5.4 kb *ApaI-NheI* fragment. For the *Lhx6* targeting vector, the 2 kb genomic fragment between the homology regions is replaced by a 4 kb cassette, containing the following: (1) the 1b, 2 and 3 coding exons flanked by loxP sites (2) the neomycin resistance gene under the control of the phosphoglycerate kinase (PGK) promoter (PGK-Neo) flanked by FRT sites. Targeting constructs were linearized and electroporated into E14Tg2A embryonic stem (ES) cells. Targeted clones were identified and analyzed in detail by Southern blotting using the 5’ and 3’ external probes indicated in Supplementary Figure 1B. Germline transmission of the mutant alleles was achieved using standard protocols. The phenotypic analysis presented was performed on animals from which the PGK-Neo cassette was removed by crossing founder *Lhx6^Fl/+^* animals with the *Tg(ACTFLPe)9205Dym* transgenic line (MGI:2448985, Rodriguez et al., 2000). Deletion was confirmed using Southern blotting analysis. For the maintenance of the *Lhx6^Fl^* colony, we performed PCR using the following primers: F:5’-<u>CTCGAGTGCTCCGTGTGTC</u>-3’, R:5’-<u>GGAGGCCCAAAGTTAGAACC</u>-3’.

### Animals

Animals were bred and housed in accordance with the United Kingdom Animals (Scientific Procedures) Act (1986). The *Tg(Nkx2-1-cre)1Wdr* (MGI:3761164, shortened here as *Nkx2.1Cre,* Kessaris et al., 2006), *Slc32a1^tm2(cre)Lowl^* (MGI:5141270, shortened here as *VGatCre,* Vong et al., 2011), *Tg(Htr3a-cre)NO152Gsat* (MGI:5435492, shortened here as *Htr3aCre,* generated by The Gene Expression Nervous System Atlas –GENSAT - Project, The Rockefeller University - New York), *Gt(ROSA)26Sor^tm14(CAG-tdTomato)Hze^* (MGI:3809524, shortened here as *Ai14,* Madisen et al., 2010). *Lhx6^tm2Vpa^* (MGI:3702518, shortened here as *Lhx6^-^*, Liodis et al., 2007) and **Lhx6^fl^** animals were maintained on a mixed background and genotyped as described previously. To generate productive crosses, *Cre-positive;Lhx6^+/-^* males were mated with *Lhx6^fl/fl^; Ai14* or *Lhx6^+/-^; Ai14* females and the resulting littermates were analyzed.

### Illumina RNA-Seq library preparation, sequencing and analysis

Illumina RNA-Seq libraries were made with 1 μg of total RNA according to the manufacturer’s protocol. The only deviation from the protocol was to use the e-gel clone well system (Invitrogen) for fragment size selection. Total RNA was extracted from the forebrain of P15 mice. Each sample was composed of two animals. Four samples for *Lhx6^+/-^* and three samples for *Lhx6^-/-^* mouse brains were collected and further analysed. 36 base pair single end sequencing was undertaken using an Illumina GA IIx DNA sequencer. FASTQ reads were trimmed with CutAdapt (http://journal.embnet.org/index.php/embnetiournal/article/view/200/479) to remove adapter contamination and low quality sequence. Reads were aligned to the Ensembl GRCm38.84 version of the mouse transcriptome using Bowtie2 (Langmead and Salzberg, 2012). Expected gene counts were extracted from the alignments using RSEM (Li and Dewey, 2011). Differential expression analysis was then perfomed using DESeq2 package (Love et al., 2014). Significance was assigned to genes with an adjusted p-value of <= 0.05. <u>Gene Ontology (GO) analysis</u>: Gene lists were taken from Broad molecular signatures database (MsigDB) (http://software.broadinstitute.org/gsea/msigdb/). Human orthologues of the differentially expressed mouse genes were identified using the Ensembl BioMart database. Significance enrichment of differentially expressed genes within MsigDB gene lists were identified using the hypergeometric test. To correct for multiple-hypothesis-testing a false-discovery rate was calculated from the resultant p-values. <u>Heatmaps</u>: The log2-fold-change expression, of each gene in each sample, was calculated relative to the mean expression of the gene across all samples. Hierarchical clustering of the samples was performed on the euclidean distances between the log2-fold-change values for each sample.

### Quantification of CGE-derived CI transcripts

For quantification of transcripts in CGE-derived CIs, P7 brains were dissociated (papain dissociation system, Worthington, Lorne) and tdT^+^ cells were isolated by flow cytometry (FACS ARIAII, Becton-Dickinson; typically, a yield of 60,000 cells). Total RNA was purified (RNeasy microKit, Qiagen) and subjected to reverse transcription (RT^2^ First Strand, Qiagen) and quantitative PCR according to manufacturer’s instructions (RT^2^ Profiler PCR plasticity pathway-focused gene expression array, Qiagen). Experiment represents results from two *Htr3aCre;Ai14;Lhx6^-/-^* and two *Htr3aCre;Ai14;Lhx6^+/-^* samples.

### Expression constructs

Full length cDNA for hM3D(Gq) from plasmid pAAV-hSyn-HA-hM3D(Gq)-IRES-mCitrine (Addgene, 50463), was cloned into a modified pCAGGS-IRES-RFP vector (a gift from Francois Guillemot - CRICK), resulting in pCAGGS-hM3D(Gq)-IRES-RFP. The pCAGGS-IRES-GFP was a gift from James Briscoe (CRICK).

### GE cell transplantations

Both medial and caudal ganglionic eminences were dissected from E14.5 *Gad1^tm1.1Tama^* (MGI:3590301, shortened here as GAD67::GFP, Tamamaki et al., 2003) heterozygote embryos, dissociated as previously described (Du et al., 2008), and the resulting cell suspension was grafted into the cortices of *Lhx6* control and mutant neonatal pups (P0-P1). One single injection has been performed into the cortex of each pup, by using the microinjector unit set-up from the VEVO injection system (VisualSonics). Each injection (69 nl) has been performed at a slow injection rate (23nL/sec) with needles according to the recommended dimensions of the manufacturer. The same needle has been used for all injections to pups of the same litter, and between each injection the needle has been inspected to verify the same cell suspension volume was injected. Grafted animals were transcardially perfused at P16, and dissected brains were processed for immunohistochemistry. Only litters containing at least one *Lhx6^-/-^* (mutant) and one *Lhx6^+/-^* (control) mouse were analysed, and values for mutants were normalized to the average number found in the control littermates, injected with the same cell suspension.

### MGE electroporation and cell transplantations

*Ex vivo* electroporation of MGE in embryonic brain slices (E14) was conducted as described previously (Stuhmer et al., 2002). Twelve hours after electroporation of a mixture of pCAGGS-hM3D(Gq)-IRES-RFP and pCAGGS-IRES-GFP plasmids, MGE regions with the strongest GFP signal were dissociated as previously described (Du et al., 2008). The resulting cell suspension was grafted into neonatal (P0-P1) cortices of wild type mice (6 injections per brain/3 per hemisphere) as described above. Cohorts of littermates grafted with the same cell suspension were divided in two groups: one group was injected intraperitoneally twice per day (every 12 hrs) with 1mg/kg CNO (Tocris Bioscience) (diluted in vehicle - 0.5% DMSO containing saline); while a control group was injected with vehicle only, from P14 until P17 (one injection only at P17). Mice where then transcardially perfused within 1 hr from the last injection, and dissected brains were processed for immunohistochemistry. Only experiments where GFP^+^ cells were identified in at least one animal from each group were analysed.

### In utero elecroporations

*In utero* electroporation was carried out using a protocol adapted from (Saito and Nakatsuji, 2001). Briefly, timed-pregnant wild type CD-1 female mice were anesthetized with a mix of oxygen-isoflurane before the abdomen was opened and the uterine horns exposed. The DNA solution was injected into the lateral ventricle of E14.5 embryos using a glass micropipette. Approximately 0.5 microliters of a solution containing Tris-HCl (10 mM), ethylenediaminetetraacetic acid (EDTA, 1 mM), Fast green dye (0.5% w/v) and a mixture (3:1 molar ratio) of pCAGGS-hM3D(Gq)-IRES-RFP and pCAGGS-IRES-GFP plasmids (total DNA concentration ≈ 1 mg/ml) was injected. Five square electric pulses (40 V, 50 ms) were passed at 1 s intervals using a square-wave electroporator (CUY21EDIT, NEPA GENE Co.). At P21 a group of mice received an intraperitoneal injection of 1mg/kg CNO (Tocris Bioscience) (diluted in vehicle - 0.5% DMSO containing saline) and were transcardially perfused and processed for immunohistochemichestry with GFP, RFP and cfos antibodies. A group of control mice did not receive CNO injections.

### Immunostaining

For immunostaining on brain sections from P16 mice or older, animals were transcardially perfused with 4% PFA and brains were post-fixed overnight (O/N). Vibratome (60 or 100 µm) sections were permeabilized with PBT [0.5% Triton X-100 in PBS (0.5% PBT)] for 1 hr at room temperature (RT), blocked in 10% FCS in PBT (0.3% Triton X-100 in PBS; 2 hrs; RT), and incubated with primary antibodies diluted in blocking solution at 4°C, O/N. After 3 washes with PBT, sections were incubated with secondary antibodies diluted in blocking solution at RT for 2 hrs, washed in PBT, and mounted using Vectashield (Vector) medium. For immunostaining on brain sections from embryos (E14.5, E16.5) or P2/P7 mouse pups, dissected brains were fixed in 4% PFA in PBS at 4°C, O/N. Cryostat sections (14 µm) were permeabilized in 0.1% Triton X-100 in PBS (0.1%PBT) for 5 min, and processed as above. The following antibodies were used: rabbit polyclonal anti-cFos (Calbiochem; 1/10.000), mouse monoclonal anti-cFOS (Santa-Cruz, sc-166940; 1/500), rabbit polyclonal anti-GFP (Invitrogen; 1/1000), rat monoclonal anti-GFP (Invitrogen; 1/1000), mouse monoclonal anti-GFP (Invitrogen; 1/1000), rabbit polyclonal anti-Lhx6 (Lavdas et al., 1999; 1/250–1000), rabbit polyclonal anti-Parvalbumin (Swant, PV25; 1/1000), goat polyclonal anti-Parvalbumin (Swant, PV213; 1/1000), rabbit anti-PH3 (Millipore, 05-636; 1/200), rabbit polyclonal anti-RFP (Abcam; 1/500), mouse monoclonal anti-Reelin (Millipore, MAB5364; 1/500), rabbit polyclonal anti-Sox6 (Abcam, AB30455; 1/4000), goat polyclonal anti-Sp8 (Santa-Cruz, sc-104661; 1/1000), rat monoclonal anti-SST (Millipore, MAB354; 1/1000), rabbit polyclonal anti-VIP (Immunostar, 20077; 1/1000). Secondary antibodies used were as follows: Alexa Fluor 488-conjugated donkey anti-mouse, anti-rat, anti-goat and anti-rabbit and Alexa Fluor 568-conjugated donkey anti-mouse, anti-rat, anti-goat and anti-rabbit (all from Invitrogen; all 1/500).

### In situ hybridization histochemistry (ISHH)

ISHH was carried out essentially as described (Schaeren-Wiemers and Gerfin-Moser, 1993). The Arc-specific and Egr1-specific riboprobes were a gift from Dr. Tahebayashi.

### EdU (5-ethynyl-2’-deoxyuridine) injection/staining

A stock solution of 10mg/ml EdU (Invitrogen, E10187) was prepared in DPBS (Life technologies, 14190-094). Pregnant females were injected intraperitoneally with 3µl/g weight of each mouse. To study the fraction of cells in the S-phase of the cell cycle, pregnant females were injected 1 hour prior to embryo harvest. Brain cryosections from E14.5 or E16.5 embryos were first processed for immunohistaining (PH3) and then EdU detection according to the manufacturer’s protocol.

### Tunel terminal deoxynucleotidyl transferase-mediated dUTP nick end-labelling) assay

In order to detect cells that are undergoing apoptotic cell death, we used the ApopTag Green *In Situ* Apoptosis Detection Kit (Invitrogen, S7111) and followed manufacturer’s instructions. Briefly, immunostained sections were fixed for 10 minutes with 4% PFA, washed with PBS and post-fixed further with an EtOH/Acetic Acid (2:1) mix. Upon equilibration with the appropriate buffer, sections were incubated with TdT enzyme mix for 1hour at 37°C. The reaction was then stopped with STOP buffer, sections were washed and slides were incubated with fluorescin conjugated anti-DIG mix in a dark humidified chamber, for 30 minutes at room temperature. Slides were then washed thoroughly with PBS and mounted with Vectashield-DAPI mounting medium.

### Image analysis

Cell counting for all post-natal stages was performed from coronal sections in the cortical region. Images were acquired using a confocal microscope (×20 magnification), taking care for all pairwise comparisons to be acquired in the same imaging session with the same acquisition settings. Cells were manually identified in columnar regions spanning the pial-white matter extent of the cortex across different bregma levels between +2 and -3 mm (as defined in Paxinos et al., 2001) or equivalent regions in the early post-natal stages shown in Fig. 2 A-H). Bregma regions were closely matched for each pairwise comparison. No obvious changes in the reported changes were detected between different bregma regions (e.g. results in Fig. 1 G-M are similar to Fig. S2). tdT^+^ cell numbers in Fig. 1 G-I, V and Fig. S2 A,D were divided by the total surface area counted (measured in ImageJ) to obtain cell densities and normalized to the average density observed for control animals in each pairwise comparison. For analysis of specific CI subpopulations (Fig. 1 J-M and Fig. S2 B,C,E,F, separate channels were acquired and markers were initially assessed independently, and were later combined to assess co-localization using the ImageJ software. Results were presented as cell densities (cells per mm^2^). Essentially similar methods were used for analysis of apoptosis of fate mapped CI populations (Fig. 2 K-L) and characterization of the MGE CI population in *Lhx6* conditional mutants (Fig. S1 I-N), except that cell numbers were normalized to the size of the tdT^+^ population. For Figs 1 G-M analysis was restricted to the primary somatosensory cortex (as defined in Paxinos et al., 2001), while for Fig. S2 countings were performed in the primary motor cortex. For Fig. 2 I, K-L and Fig. S1 G-M countings were performed for at least 4 different matched bregma levels and measurements were averaged to obtain cell densities for each animal. For analysis of survival of grafted GAD67::GFP^+^ CIs (Fig. 3 G), cells present in the cortex in all sections were counted. Numbers for each litter were normalized to the average number found in the *Lhx6^+/-^* animals, a minimum of 500 cells were counted per brain. For analysis of survival of grafted CIs transfected with hM3D(Gq) expressing constructs (Fig. 5 J) GFP^+^ cells present in the cortex in all sections were counted and assessed for RFP expression. Numbers of double positive cells (RFP^+^GFP^+^) were expressed as a fraction of total GFP^+^ population for each brain, a minimum of 150 cells were counted per brain. In the graph lines connect markers corresponding to animals grafted with the same cell suspension and given different treatments (vehicle or CNO). For analyses of soma size (Fig. S5), an oval shaped region of interest (ROI) was drawn just inside the soma region of cells that were judged to be in focus. The area size of the ROIs was measured. Area sizes were measured from at least three different brains in matched bregma levels. For analysis of nuclear cfos expression (Fig. S6) images were acquired at 40X magnification. The cfos channel was analysed independently, by drawing oval shaped ROIs inside the nuclear region of all nuclei present in the image, at the Z slice with highest fluorescent intensity. Average fluorescent intensity was measured for all ROIs in both the cfos channel and the RFP channel. Measurements in the cfos channel were divided in two groups based on RFP intensity. All cfos intensity measurements were normalized to the median intensity of the group with RFP expression below threshold. All imaging and analysis were done by an experimenter blind to the experimental condition. All values are presented as mean ± standard error of the mean. All results presented in Figures are presented in the Supplementary Table S1, accompanied by the reference to the Figure panel where they appear, and the statistical test used to assess significance.

